# 4 Phenylbutyrate Plus Gene augmentation: A Dual Therapy To Rescue of SLC6A1 Variant Associated Developmental And Epileptic Encephalopathy

**DOI:** 10.64898/2026.06.05.730491

**Authors:** Aiden James Delahanty, Kaitlin James, Emma Grace, Ziasng Debbie Song, Juexin Wang, Melissa Bassette, Jing-Qiong Kang

## Abstract

**Background:** Pathogenic variants in SLC6A1, which encodes the γ-aminobutyric acid (GABA) transporter GAT-1, cause developmental and epileptic encephalopathies (DEEs) through reduced GABA uptake, impaired transporter trafficking, and functional haploinsufficiency. 4-Phenylbutyrate (PBA) is a clinically available small molecule with chemical-chaperone and histone-deacetylase-inhibitor activities that can rescue misfolded GABAergic proteins, but variant-level rescue data are needed to guide precision treatment.

**Methods:** We report a newly identified de novo SLC6A1 missense variant, p.Ala305Val (A305V), in a patient with myoclonic-atonic epilepsy and a developmental and epileptic encephalopathy phenotype. A305V was compared with the residue-matched comparator p.Ala305Thr (A305T). Variant effects were evaluated by (i) protein-structure prediction across nine stability-prediction algorithms using the cryo-EM-derived human GAT-1 template (PDB 7Y7W); (ii) 3H-GABA uptake assays in HEK293T cells and in human iPSC-derived astrocytes and cortical neurons; (iii) live-cell confocal microscopy of ER colocalization; (iv) pharmacologic rescue with PBA, TUDCA and salubrinal (v) and GAT-1 cDNA gene-augmentation, alone and in combination with PBA.

**Results:** AI-based stability predictors uniformly indicated destabilization of GAT-1(A305V) and GAT-1(A305T). A305V reduced ^3^H-GABA uptake across HEK293T, astrocyte, and neurons. The mutant transporter accumulated within the endoplasmic reticulum (ER), with ER colocalization rising from approximately 30% in wildtype to ∼80% in A305V; PBA reduced ER retention to approximately ∼40% and restored total GAT-1 fluorescence toward wildtype levels. Pharmacochaperones (PBA, TUDCA) restored GABA uptake for the mutant transporters. Wildtype GAT-1 gene augmentation improved mutant GAT-1 uptake and combined PBA-plus-augmentation produced rescue greater than either intervention alone in the available dose-response ranges.

**Conclusions:** SLC6A1 A305V is a trafficking-impaired, loss-of-function GAT-1 variant whose dysfunction is tractable to two convergent therapeutic axes: pharmacologic correction of folding and trafficking, and augmentation of functional transporter dose. These findings support a two-pronged precision-medicine framework for SLC6A1-related DEEs in which PBA increased the transporter function augmented by increased gene therapy.

**Significance of the study:** This work links the patient-derived SLC6A1 A305V variant to a defined molecular mechanism—GAT-1 destabilization, ER retention, and reduced GABA uptake—and demonstrates that the deficit is reversible by two independent interventions that converge at the same downstream endpoint of functional surface transporter. Because PBA is already clinically deployable and GAT-1 cDNA augmentation models a future viral or non-viral gene therapy, the combined-rescue logic provides a falsifiable path for precision medicine in SLC6A1-related DEEs: chemical chaperoning corrects the folding bottleneck while transporter augmentation increases the pool available for rescue.

## Introduction

γ-Aminobutyric acid (GABA) transporter 1 (GAT-1), encoded by SLC6A1, is the principal neuronal and astrocytic transporter that clears GABA from the synaptic cleft and extrasynaptic compartments in the mammalian central nervous system (Roth and Draguhn 2012; Eulenburg and Gomeza 2010). By terminating GABAergic neurotransmission and shaping tonic inhibition, GAT-1 helps maintain the excitation–inhibition balance whose loss underlies generalized and developmental epilepsies. Pathogenic SLC6A1 variants disrupt this homeostasis and cause a spectrum of developmental and epileptic encephalopathies (DEEs) typically characterized by myoclonic-atonic seizures, absence seizures, atypical absences, global developmental delay, autism-spectrum features, attention-deficit/hyperactivity symptoms, and sleep or behavioral disturbances (Carvill et al. 2015; Johannesen et al. 2018; Stefanski et al. 2023).

We have previously characterized the mutant GAT-1 function and trafficking in various cell and mouse models (Mermer et al., 2021, Nwosu et al., 2022). Work from our laboratory and others has shown that the majority of pathogenic SLC6A1 missense variants do not simply abolish substrate translocation. Rather, they destabilize the folded transporter, lead to glycosylation arrest and endoplasmic reticulum (ER) retention, and produce a functional haploinsufficiency in which mutant protein fails to reach the plasma membrane in a transport-competent state (Cai et al. 2019; Wang et al. 2020; Mermer et al. 2021; Mermer et al. 2022; Shen et al. 2024). This recurring feature—a trafficking-linked loss of function rather than a pure channelopathy—reframes SLC6A1-related DEE as, in part, a proteostasis disease and points toward therapeutic strategies that target folding and forward trafficking rather than channel pharmacology alone.

4-Phenylbutyrate (PBA), an FDA-approved small molecule used clinically for urea-cycle disorders, has chemical-chaperone activity, low-grade HDAC-inhibitory activity, and ER-stress-modulating activity. Importantly, we have identified 4 phenylbutyrate, can restore the GABA uptake and mitigate seizures in the cell and mouse models (Nwosu et al. 2022) as well as human patients bearing the SLC6A1 variants (Grinspan et al, 2024). In preclinical SLC6A1 and GABAₐ-receptor models, PBA improves protein folding, reduces ER stress, increases surface expression, restores GABA uptake or receptor function, and mitigates seizure phenotypes (Nwosu et al. 2022; Shen et al. 2023). An ongoing investigator-initiated clinical trial of glycerol phenylbutyrate (Ravicti) in SLC6A1 and STXBP1 (NCT04937062) provides an unusually strong translational foothold for a rare-disease program. However, both clinical observation and preclinical work indicate that the magnitude and mechanism of PBA rescue are variant-dependent, and a variant-by-variant characterization remains the rate-limiting step in precision deployment.

Computational protein-stability prediction is increasingly used to triage variants of uncertain significance and to forecast the structural consequences of missense substitutions (Rodrigues et al. 2018; Pires et al. 2014). When integrated with experimental validation, stability prediction does not replace biology; it focuses on biology. For a transmembrane transporter such as GAT-1, where structural data are now available at near-atomic resolution (Zhu et al. 2023), in-silico modeling provides a falsifiable expectation about whether a specific residue substitution should disrupt folding or transport, and that expectation can be tested directly with uptake and imaging assays.

Here we report the functional, structural, and rescue characterization of SLC6A1 p.Ala305Val (A305V), a newly identified de novo missense variant associated with developmental and epileptic encephalopathy. We have previously elucidated the mechanism of pharmacochaperoning and HDAC inhibitor as well as the effect of by genetic upregulating the folding machinery like Bip and Calnexin. The central translational hypothesis is that chemical chaperoning underlying transporter upregulation. Here we evaluate the convergent therapeutic axes: pharmacologic correction of protein misfolding (PBA, TUDCA and salubrinal), and augmentation of the wildtype allele with or without the treatment of PBA. Considering the ongoing gene therapy as a promising treatment option for many epilepsy and DEEs, the study provides proof of concept understanding for two different approaches, small molecules and gene therapy as complementary rather than redundant for trafficking-impaired SLC6A1 variants. PBA can optimize and de-risk gene therapy by maximally leveraging the existing wildtype allele and rescuable mutant allele.

## Methods

### Patient ascertainment, clinical evaluation, and genetic diagnosis

The proband and available family members were evaluated at Vanderbilt University Medical Center. Written informed consent for the use of clinical and genetic information was obtained from the parent(s) or legal guardian(s) prior to inclusion. Abstracted clinical data included age of seizure onset, seizure semiology and frequency, developmental milestones, antiseizure medication exposure and response, video-EEG findings, and neurodevelopmental milestones and comorbidities such as autism-spectrum disorder and attention-deficit/hyperactivity symptoms were assessed using standard instruments. Saliva samples were submitted for CLIA-approved clinical genetic testing on the Invitae Comprehensive Epilepsy Gene Panel (Invitae, San Francisco, CA, USA). The SLC6A1 p.Ala305Val variant was identified *de novo*.

### Protein-structure prediction and machine-learning analyses

Stability and flexibility changes induced by A305V and A305T were predicted from the cryo-EM–derived human GAT-1 template (PDB 7Y7W) (Zhu et al. 2023), which was selected for its 2.9-Å resolution and complete coverage of the Ala305 site. Point mutations were introduced in silico, and ΔΔG predictions were obtained from DynaMut2 (Rodrigues et al. 2021), mCSM (Pires et al. 2014), SDM (Worth et al. 2011), DUET (Pires et al. 2014), I-Mutant, MUpro (SVM and neural-network variants), INPS-MD (Savojardo et al. 2016), and MAESTROweb (Laimer et al. 2016). The same structural template was used across models for consistency, and effects were interpreted in the context of side-chain polarity, hydrophobic packing, steric fit, and transmembrane architecture. Negative ΔΔG values indicate predicted destabilization.

### Expression constructs and site-directed mutagenesis

Plasmids encoding human GAT-1 tagged with enhanced yellow fluorescent protein (EYFP) were constructed in the pCMV mammalian expression vector as previously described (Cai et al. 2019; Wang et al. 2020). The A305V and A305T missense substitutions were introduced into the wildtype GAT-1 coding sequence using the QuikChange Site-Directed Mutagenesis Kit (Agilent), transformed into DH5α competent cells, plated under antibiotic selection, and verified by Sanger sequencing of single colonies. Endotoxin-controlled plasmid stocks were prepared with the QIAGEN Maxiprep kit and stored at –20°C. For heterozygous modeling, wildtype and mutant constructs were co-expressed at equal mass ratios; for homozygous or mutant-only modeling, mutant cDNA alone was transfected.

### Polyethylenimine (PEI) transfection

Standard transfection protocols were performed using human embryonic kidney 293T (HEK293T) cells (Masur D, 2013). 24 hours before transfection HEK293T cells were split equally into plates. For radiolabeling GABA uptake,1µg of the cDNAs with PEI at a ratio of 1:2.5µl was transfected in 35mm dish or in 60 mm dish for western blot. The cDNAs were combined with Dulbecco modified Eagle medium (DMEM) and a PEI/DMEM mixture. For total protein expression, 3µg cDNAs were used for transfection. Transfected HEK293T cells were incubated for 48 hours. After incubation, proteins were harvested as described below.

### Cell culture, iPSC differentiation for neurons and astrocytes, and transfection

HEK293T cells were maintained in Dulbecco’s Modified Eagle’s Medium (DMEM) with 10% fetal bovine serum and 1% penicillin/streptomycin. Transfections used polyethylenimine (PEI) at a 1:2.5 mass ratio of cDNA to PEI, with 1 μg cDNA per 35-mm dish for GABA-uptake assays and 3 μg cDNA per dish for total-protein analysis. Transfections were incubated for 48 h prior to assay.

Human induced pluripotent stem cells (iPSCs; Thermo Fisher A18945) were maintained and differentiated into neural progenitor cells (NPCs), astrocytes, and cortical inhibitory neurons following our previously validated protocol (Mermer et al. 2021). NPC induction used the STEMdiff SMADi Neural Induction Kit (STEMCELL Technologies). Astrocytes were differentiated from P1 NPCs in Astrocyte Medium (ScienCell) for 25–30 days; medium was refreshed every other day, and cells were passaged at ≈70% confluence. Astrocyte identity was confirmed by S100β and glial fibrillary acidic protein (GFAP) immunostaining, with >95% of cells adopting astrocytic morphology and ≈80% staining GFAP-positive, consistent with prior reports. Cortical inhibitory neurons were differentiated from passage-2 NPCs at day 10 of induction, harvested at differentiation days 60–65, and validated by NeuN, DLX, synapsin, and synaptophysin immunostaining. Astrocytes and neurons were transfected with 1 μg cDNA per 35-mm dish using Lipofectamine.

### ^3^H-GABA uptake assay

Transporter function was measured using a modified ^3^H-GABA uptake assay (Keynan et al. 1992; Cai et al. 2019). HEK293T cells were seeded at 1.5 × 10⁵ cells per 35-mm dish three days before assay and transfected with 1 μg of wildtype or mutant GAT-1 cDNA at 24 h after seeding. Uptake was performed 48 h after transfection. Cells were preincubated with uptake buffer for 15 min, then exposed to 1 μCi/mL 3H-GABA and 10 μM unlabeled GABA at room temperature for 30 min. After washing, cells were lysed with 0.25 N NaOH for 1 h, neutralized with glacial acetic acid, and counted on a liquid-scintillation counter (QuantaSmart). GABA flux (pmol/μg protein/min) was averaged from technical triplicates per condition; untransfected wells served as background and were subtracted. Mutant uptake was normalized to wildtype within each experiment, with wildtype defined as 100%. The GAT-1–selective inhibitor Cl-966 (100 μM) and the GAT-3–selective inhibitor SNAP5114 (30 μM, astrocytes only) were used as pharmacologic specificity controls.

### Pharmacologic and gene augmentation rescue

Cells expressing wildtype or mutant GAT-1 were treated with vehicle for control and PBA (2 mM, 24 h), TUDCA (100 μM, 24 h) or salubrinal (15 μM, 24 h) prior to GABA-uptake or imaging endpoints. For genetic augmentation with cDNAs, GAT-1 constructs were co-transfected with empty PcDNA control at desired mass ratios to normalize the total transfected cDNA amount when necessary.

### Gene augmentation and combined rescue

GAT-1 cDNA augmentation was modeled by titrating wildtype GAT-1 plasmid (0.125, 0.25, or 0.5 μg) into mutant-expressing astrocyte cultures while holding total transfected DNA constant with carrier PcDNA. GAT-1(A305V) and GAT-1 (S295L) were tested in parallel as trafficking-impaired comparator variants. For combined rescue, GAT-1(A305V) and GAT-1(S295L) expressing cells received fixed GAT-1 augmentation in the presence of increasing PBA concentrations (0.5, 1, or 2 mM, 24 h). The combination is described throughout as gene augmentation rather than gene therapy because *in vivo* viral delivery in animal was not performed in this study.

### Live-cell confocal microscopy and ER colocalization

Cells were plated on poly-D-lysine–coated, glass-bottom dishes at 1–2 × 10⁵ cells/dish and co-transfected with 0.5 μg of wildtype or mutant GAT-1 tagged with enhanced yellow fluorescent protein (EYFP) (GAT-1^YFP^) and 0.5 μg of enhanced yellow fluorescent protein (ECFP)-tagged ER marker (ER^CFP^) with using PEI. Live-cell confocal imaging was performed 48 h post-transfection on a Zeiss LSM 510 inverted laser-scanning microscope with a 63×/1.4 NA oil-immersion objective and multi-track excitation at 458 nm (ECFP) and 514 nm (EYFP). Single confocal sections were averaged from eight frames to reduce noise unless otherwise noted. ER colocalization was quantified in MetaMorph and ImageJ as the percentage of GAT-1^YFP^ fluorescence overlapping with the ER marker, and total GAT-1 fluorescence intensity was measured in parallel.

### Western blot of GAT-1 expression

Transfected cells were washed in PBS, lysed in RIPA buffer (20 mM Tris, 20 mM EGTA, 1 mM DTT, 1 mM benzamidine, 0.01 mM PMSF, 0.005 g/mL leupeptin, 0.005 g/mL pepstatin) for 30 min at 4°C, and resolved by SDS-PAGE. Membranes were probed with rabbit anti–GAT-1 (Alomone AGT-001 or Synaptic Systems 274102; 1:200) and quantified with Odyssey. GAT-1 ran as three bands at 108, 96, and 90 kDa, corresponding to mature glycosylated, partially glycosylated, and immature core-glycosylated forms, respectively, consistent with our prior reports (Cai et al. 2019). Mature/immature band-intensity ratios were used as a maturation readout.

### Statistical analysis

Numerical data were expressed as mean ± SEM. Proteins were quantified by Odyssey software and data were normalized to loading controls and then to wildtype GAT-1, which was arbitrarily taken as 1 in each experiment. The radioactivity of GABA uptake was measured in a liquid scintillator with QuantaSmart. The flux of GABA (pmol/µg/min) in the wildtype GAT-1 samples was arbitrarily taken as 100% each experiment. The fluorescence intensities from confocal microscopy experiments were determined using MetaMorph imaging software and the measurements were carried out in ImageJ as modified from previous description^2,14^. For statistical significance, we used one-way analysis of variance (ANOVA) with Newman-Keuls test or Student’s unpaired *t*-test. In some cases, one sample *t*-test was performed (GraphPad Prism, La Jolla, CA), and statistical significance was taken as p < 0.05.

## Results

### A novel variant SLC6A1(A305V) and a dual therapy for rescuing SLC6A1

Previous studies have identified mutations in *SLC6A1* associated with a spectrum of epilepsy syndromes and neurodevelopmental disorders^1,2,8,15,16^. We have previously studied *SLC6A1* mutations associated with a wide spectrum of disease phenotypes including MAE, Lennox-Gastaut syndrome, autism, ADHD, neurodevelopmental delay due to various degrees of loss of GAT-1 function and genetic background of affected individuals. Most assayed mutations are *de novo* ((Figure 1A). Here we identified 914C to T mutation in *SLC6A1* resulting in a change of a conserved residue Alanine 305 to Valine (A305V) (Figure 1A) in GAT-1 protein (Figure 1B). This variant of unknown significance was not identified in parents, suggesting de novo. The mutant amino acid of Alanine is conserved across species, suggesting the variation of the amino acid may have significant impact on the transporter protein conformation and function.

**Figure 1.**
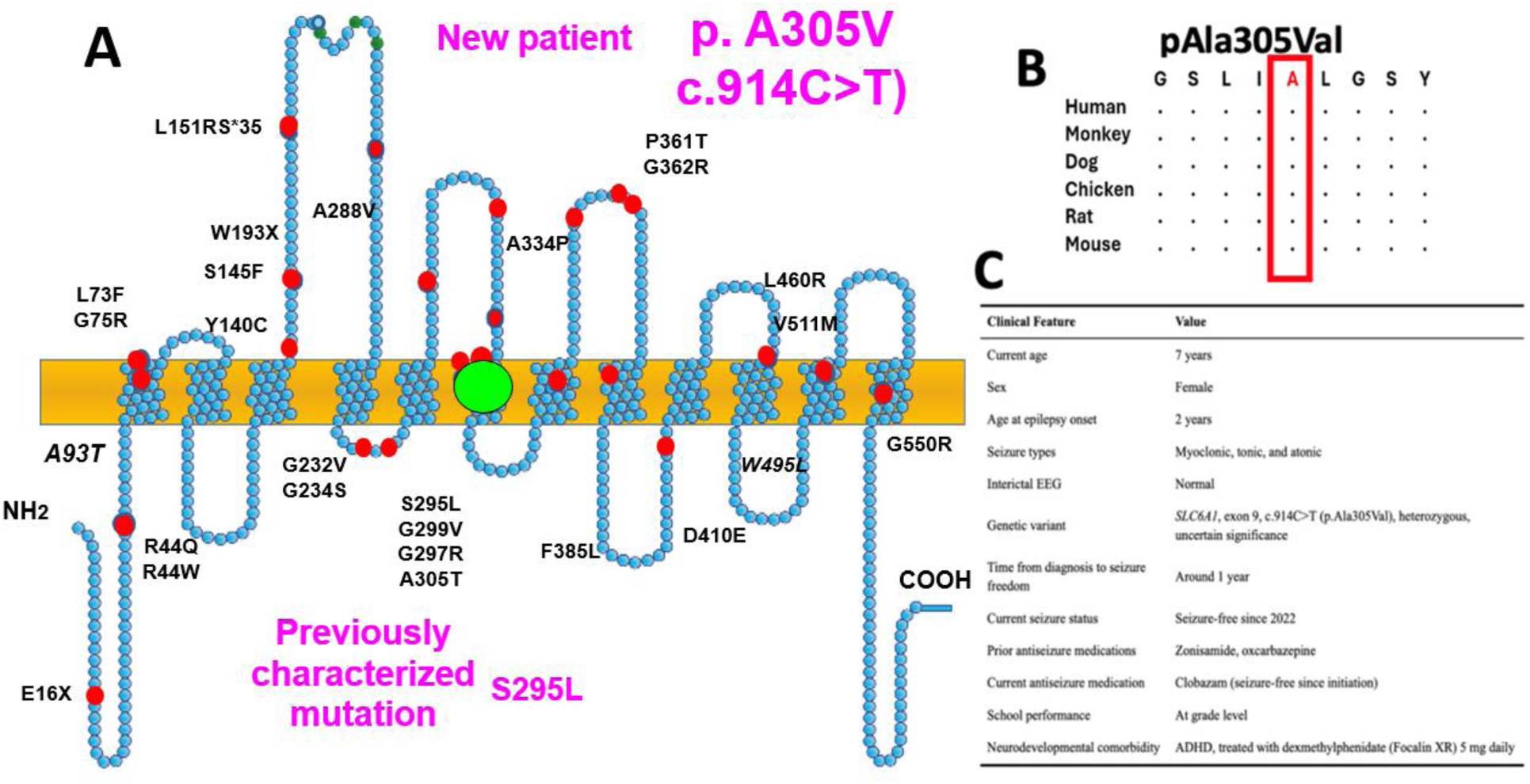
The pedigree of a novel *SLC6A1* mutation GAT1(A305V) associated with developmental and epileptic encephalopathies and experimental design of the study. **(A).** Schematic representation of GAT-1 protein topology and locations of GAT-1 variants previously identified in patients associated with developmental and epileptic encephalopathies (DEEs). It is predicted that GAT-1 contains 12 transmembrane domains. A305V is located at the 4th intracellular loop of the GAT-1 protein. The positions of variants are based on the published LeuT crystal structure. We included GAT-1(A305V) associated with other DEEs as reference Pedigree and the genotype. 914C>T>A variation resulting in GAT-1(A305V) was found de novo, in a two year-old girl but not in the mother and father. (**B**). Amino acid sequence homology shows that Alanine (A) at residue 305V is highly conserved in GAT-1 in humans (Accession NO.NP_003033.3) and across species as shown in red boxed region. (**C**). Clinical information of the patient at 7 years old.

The 7-year-old girl was born normal otherwise until onset of seizures. The first seizure onset was 2 years old and the seizure type was myoclonic, tonic and atonic. The child was treated with zonisamide, oxcarbazepine and clobazam and achieved seizure free since clobazam. The child performs at grade level with treatment for ADHD (Focalin (dexmethylphenidate) 5mg daily) (Figure 1 C).

### Computational stability prediction across nine algorithms supports destabilization at Ala305, which was validated in cells

To establish a structural expectation for the experimental program, A305V and A305T were modeled in silico against the 2.9-Å cryo-EM human GAT-1 structure (PDB 7Y7W). DynaMut2 indicated that introducing a polar threonine hydroxyl into a predominantly hydrophobic transmembrane pocket creates polarity mismatch and local steric strain at A305T, while the valine substitution in A305V increases hydrophobic bulk and creates steric hindrance in a confined packing environment (Figure 2A). Across the panel of nine stability predictors (DynaMut2, mCSM, SDM, DUET, I-Mutant, MUpro-SVM, MUpro-NN, INPS-MD, MAESTROweb), the direction of effect favored destabilization for both substitutions, with mean ΔΔG of approximately –1.4 kcal/mol for A305T and –0.5 kcal/mol for A305V; DUET predicted the largest single effect for A305T (–3.4 kcal/mol) (Figure 2B–C). The convergent prediction across mechanistically distinct algorithms provided a falsifiable expectation that experimental readouts of folding, trafficking, and transport should show comparable loss-of-function for the two substitutions.

**Figure 2.**
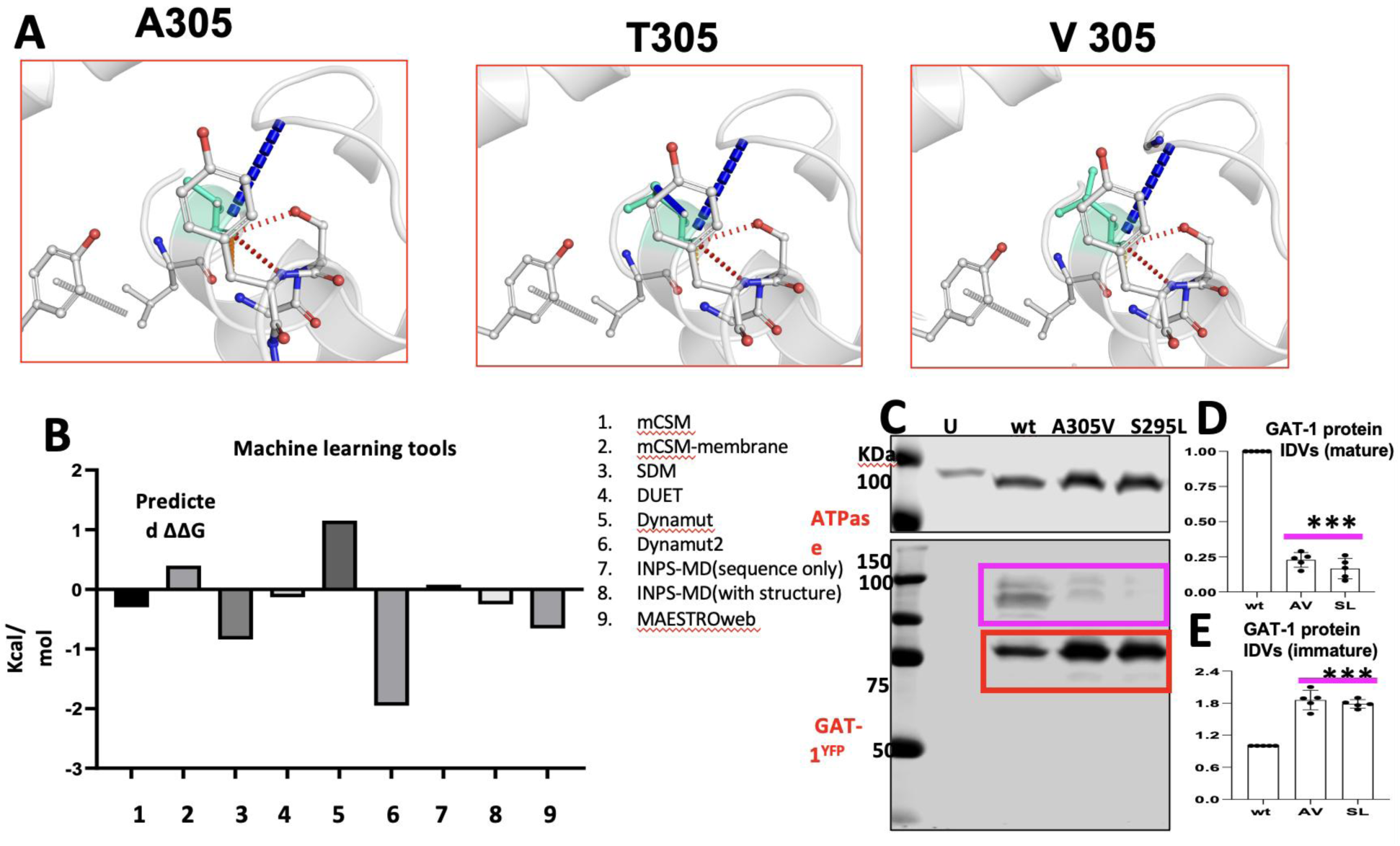
Artificial intelligence tools predict the mutant GAT-1(A305V) protein stability. (**A**). DynaMut2-predicted local environments around residue 305V (yellow arrow) in the wildtype (WT) GAT-1 and the GAT-1(A305V) mutants, using the human GAT-1 crystal structure (PDB: 7Y7W). Hydrophobic interactions are shown in green dashed lines; polar interactions (e.g., hydrogen bonds) are shown in orange dashed lines. **(B)** ΔΔG stability predictions for A305V (B) across multiple computational tools. Negative values indicate destabilization. Methods include DynaMut2 (Dyn), mCSM, SDM, DUET, I-Mutant, MUpro (SVM and NN), INPS, and MAESTROweb (MAE). (**C-E**). HEK293T cells were transfected with the wildtype GAT-1^HA^ transfected with wildtype GAT-1^YFP^; GAT-1(A305V)^YFP^ or with GAT-1(S295L)^YFP^ for 48 hours. The total cell lysates were analyzed with SDS-PAGE. The membrane was immunoblotted with a mouse monoclonal GFP antibody (C). The total protein IDVs of the mature GAT-1 in the purple box (D) or the immature GAT-1 in the red box (E) were measured. In D, E, ***p < 0.001 vs wt, n=5 gels from 5 different transfections. One-way analysis of variance (ANOVA) and Turkey post hoc test for multiple comparisons.

As shown in Figure 2A, DynaMut2 modeling suggested that introducing a threonine at position 305 places a polar hydroxyl group into a predominantly hydrophobic transmembrane pocket, leading to polarity clashes and steric strain. The valine substitution, by contrast, increases hydrophobic bulk, which can enhance local packing but also creates steric hindrance in a confined space. Across all stability prediction tools (Figure 2B), negative ΔΔG values were observed for both variants, with an average of –1.4 kcal/mol for A305T and –0.5 kcal/mol for A305V. While the magnitude of ΔΔG varied between tools, all computational tools predicted destabilization of the mutant protein, with DUET estimating the largest effect for A305T (–3.4 kcal/mol). This consistent pattern across algorithms indicates a robust destabilization signal in the mutant GAT-1.

The reduced protein stability was consistent with our biochemical assay. In the cells expressing the wildtype or the mutant GAT-1^YFP^, the GAT-1(A305V) protein at the ∼108 KDa in the purple boxed region representing the mature forms of GAT-1 was reduced compared to the wildtype (1 vs 0.228±0.023) but similar to GAT-1(S295L) (0.16±0.032), which has been identified to cause loss of GABA function due to reduced functional GAT-1 protein at the total and the cell surface levels (Figure 3 C-D). By contrast, the lower band at ∼90 KDa representing the immature form in the red boxed region was higher in the mutant GAT-1(A305V) (1.858 ±0.08) and GAT-1(S295L) (1.785 ±0.0357) compared with the wildtype (=1) (Figure 3E).

**Figure 3.**
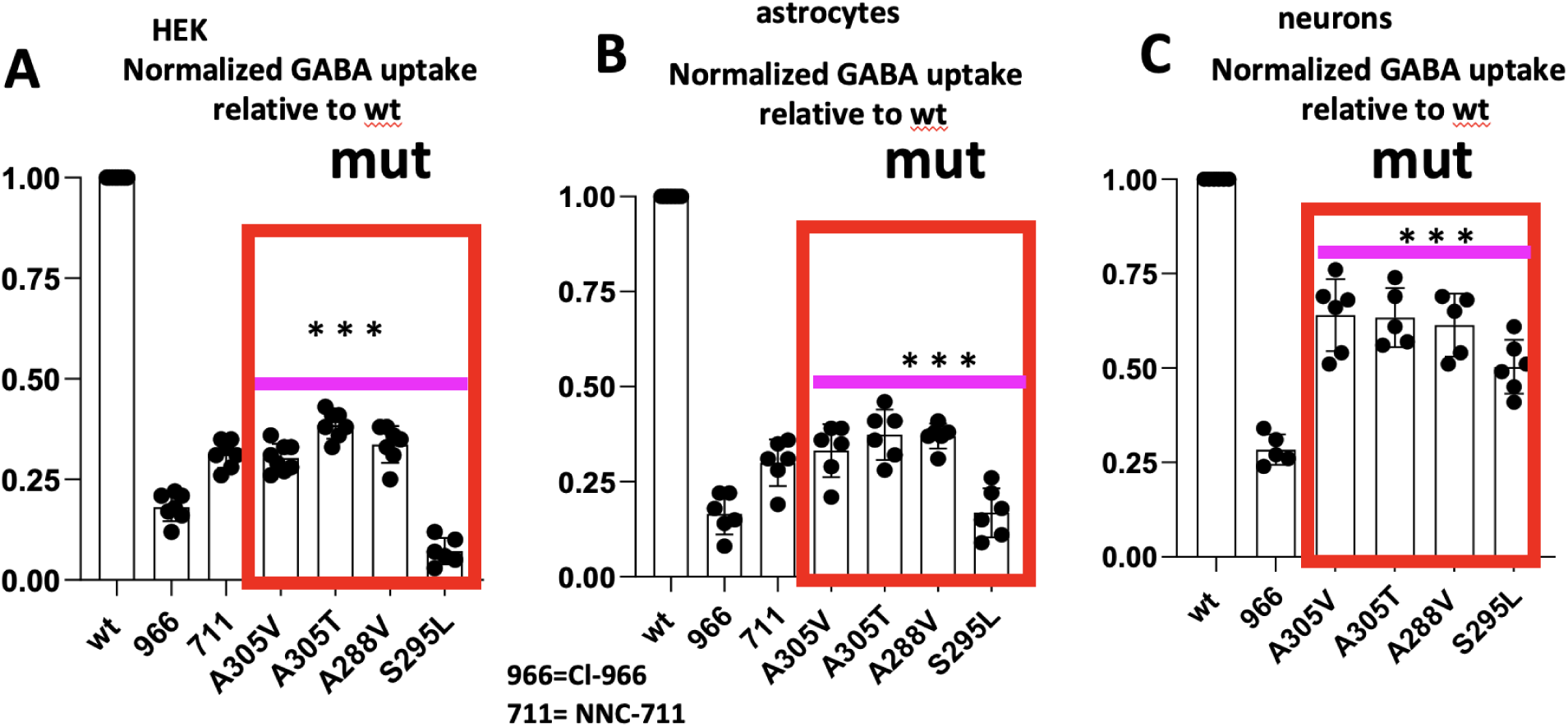
The GABA uptake function of the mutant GAT-1(A305V) and adjacent mutations across cell types including human neurons and astrocytes. **A.** HEK293T cells under passages 18 were transfected with wildtype GAT-1^YFP^ (wt), or the mutant GAT-1(A305V)^YFP^ cDNAs (0.5µg per a 35mm^2^ dish) for 48 hrs before ^3^H radioactive GABA uptake assay. **B, C**. Human iPSC derived astrocytes (**B**) and cortical neurons (**C**) were transfected with the wildtype, the mutant GAT-1(A305V) or mutant GAT-1(A305V), GAT-1(A305T) and GAT-1(S295L) cDNAs at day 27^th^ after differentiation for astrocytes and 2 months after differentiation for neurons in culture. ^3^H radioactive GABA uptake assay was performed after 2 days of transfection for astrocytes and 5 days for neurons. GABA flux was measured after 30 min transport at room temperature. The influx of GABA, expressed in pmol/µg protein/min, was averaged from duplicates of two 35-mm petri dishes for each condition and for each transfection (Carvill et al., 2015). The average counting was taken as n = 1. The untransfected condition was taken as baseline flux, which was subtracted from both the wildtype and the mutant conditions across cell types (**A**). The pmol/µg protein/min in the mutant was then normalized to the wildtype from each experiment, which was arbitrarily set as 100%. (***p < 0.001 vs. wt P<0.001; n=5-8 different transfections). The GAT-1 selective inhibitor Cl-966 (100µm) was applied 30 min across cell types while the GAT-3 inhibitor (SNAP5114 (30µM)) was only applied to astrocytes (Cl/SNAP) during GABA flux. One-way analysis of variance (ANOVA) and Turkey post hoc test for multiple comparisons.

### The A305V and A305T mutations reduce 3H-GABA uptake across HEK293T cells, human astrocytes, and human cortical neurons

In HEK293T cells, the A305V variant reduced ^3^H-GABA uptake to approximately 30% of wildtype GAT-1 function, while A305T reduced uptake to approximately 39%, similar to the uptake in cells expressing the wildtype but treated with by the GAT-1 inhibitor Cl-966 (100 μM), suggesting GAT-1 blockade (Figure 3A). In human iPSC-derived astrocytes, A305V reduced uptake to approximately 33% of wildtype and A305T to approximately 37% (Figure 3B); the GAT-3–selective inhibitor SNAP5114 was used to block GAT-3 as previously described (Mermer et al., 2022). In human iPSC-derived cortical inhibitory neurons, A305V reduced uptake to approximately 64% and A305T to approximately 61% (Figure 3C). In all three cell types, the measurements in the mutant transporter were then normalized to the cells expressing the wildtype transporter which was taken as 100% (wt = 100% vs 31 % for A305V and 37 % for A305T) (Figure 3A), human cortical astrocytes (wt = 100% vs 33 % for A305V and 37% for A305T) (Figure 3B) and neurons (wt = 100% vs 64% for A305V and 63 % for A305T) (Figure 3C). The GABA transport activity for all mutations was lower than the activity of wildtype GAT-1 and was comparable to the functionality of the wildtype transporter treated with GAT-1 inhibitors Cl-966 (100 µM). This is consistent across different cell types. In combination with the previous findings, this suggests reduced GABA uptake function is a major etiology underlying SLC6A1 variants.

### GAT-1(A305V) protein had increased endoplasmic reticulum retention while PBA can reduce the ER-bound GAT-1 protein and increased the total GAT-1 protein

Live-cell confocal microscopy demonstrated that the A305V transporter was retained intracellularly and colocalized strongly with an ER marker (ER-CFP) (4A). Approximately ∼30% of wildtype GAT-1^YFP^ fluorescence overlapped with the ER marker, compared with approximately 76% for A305V; PBA treatment reduced A305V/ER overlap to approximately 36%, restoring an expression pattern indistinguishable from wildtype GAT-1 (Figure 4A–B). Total GAT-1 fluorescence was also lower in A305V cells (approximately 25 vs. 49.7 arbitrary units in wildtype) and was restored by PBA to approximately 47.53 arbitrary units (4C). These results establish that the A305V phenotype is a folding-and-trafficking deficiency rather than a substrate-binding or kinetic deficiency, and that PBA can rescue protein folding and trafficking. Together with previous findings (Mermer at al., 2021; Nwosu et al. 2022), this suggests that increased ER retention of mutant protein and retarded wildtype protein folding due to misfolding and glycosylation arrest is a common phenomenon for mutation across GAT-1 and GABA_A_ receptor mutations.

**Figure 4.**
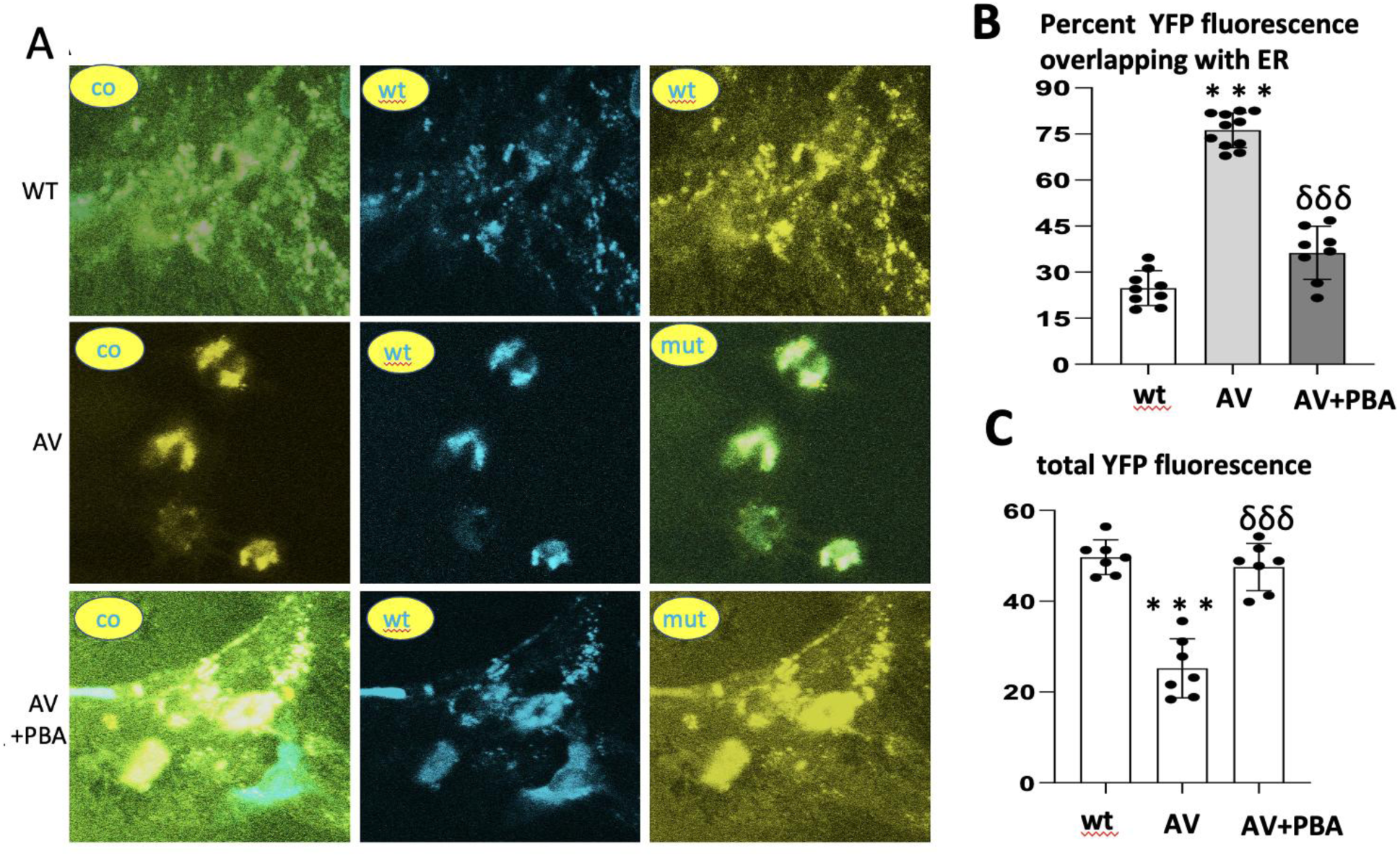
The mutant GABA transporters were retained inside the endoplasmic reticulum but were relieved by 4 phenylbutyrate in the human astrocytes. (**A**) Human astrocytes were transfected with wildtype GAT-1^YFP^ (wt) or the mutant GAT-1(A305V)^YFP^ with the pECFP-ER marker (ER^CFP^) at 1:1 ratio (1 µg:0.5µg cDNAs) for 48 hrs. The cells were treated with vehicle or 4 phenylbutyrate (2 mM) for 24 hrs before confocal microscopy. Live cells were examined under a confocal microscopy with excitation at 458 nm for CFP, 514 nm for YFP. All images were single confocal sections averaged from 8 times to reduce noise, except when otherwise specified. (**B**) The GAT-1^YFP^ fluorescence overlapping with ER^CFP^ fluorescence (**B**) or the total GAT-1^YFP^ fluorescence (**C**) was quantified by Metamorph with colocalization percentage. (**C**). PBA stands for 4 phenylbutyrate (2 mM) treated for 24 hrs. (***p < 0.001 A305VT vs. wt; &&& p < 0.001 A305V+PBA N=7 representative fields from 6 different transfections in 6 different batches of cells. One-way analysis of variance (ANOVA) and Newman-Keuls test was used to determine significance compared to the wt condition and between A305V and A305V/PBA. Values were expressed as mean ± S.E.M).

### Pharmacological chaperones and ER stress relievers increased the GABA uptake across cell types

We have demonstrated that a major mechanism for PBA to rescue SLC6A1 variants is pharmacochaperoning (Deleeuw et al. 2026). We also demonstrated that PBA can reduce ER stress in the Gabrg2^+/Q390X^ mice of Dravet syndrome (Shen et al., 2024). To dissect the mechanism by which PBA restores GABA uptake, A305V-expressing HEK293T cells were treated with PBA, the bile-acid pharmacochaperone TUDCA and the eIF2α phosphatase inhibitor Salubrinal in HEK293T cells (Figure 5A), human astrocytes (Figure 5B) and human neurons (Figure 5C). Cl-966 (50 μM) was applied as negative control. Salubrinal slows down protein translation and prevents the accumulation of misfolded proteins, thus reducing Er stress. In HEK 293T cells, the GABA uptake of the heterozygous A305V was 0.59 of the wildtype; was increased to 1.01 with PBA; to 0.815 with TUDCA and 0.82 with Salubrinal. However, compared with TUDCA and Salubrinal, PBA treatment had the best efficacy. This is consistent with our previous study in a large cohort of 32 SLC6A1 variants (DeLeeuw et al, 2026). Similarly, in both astrocytes and neurons, the GABA uptake of the heterozygous A305V was 0.66 of the wildtype; was increased to 1.04 with PBA in astrocytes and 1.03 in neurons; to 0.88 in astrocytes and 0.92 in neurons with TUDCA and 0.88 in astrocytes and 0.89 in neurons with Salubrinal. Across cell types, compared with TUDCA and Salubrinal, PBA treatment had the best efficacy.

**Figure 5.**
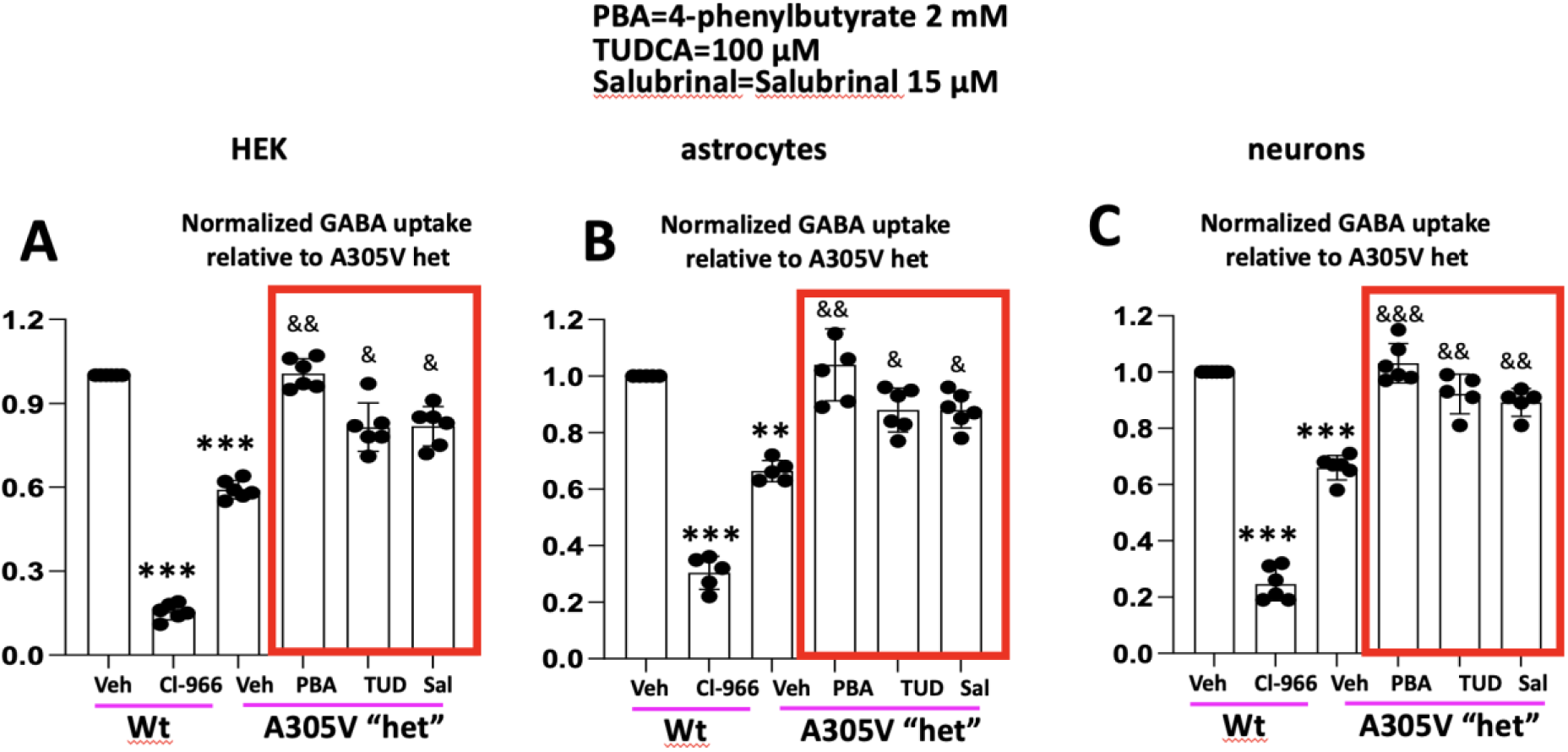
The mutant GAT-1(A305V) mutant transporters were rescued by pharmacological compounds. (**A, B**) HEK293T cells were transfected with wildtype GAT-1^YFP^ or with the mixture of the wildtype and the mutant GAT-1 (A305V) (0.25µg:0.25µg cDNAs) “heterozygously” (**B**). The cells were then treated with vehicle (Veh), 4 phenylbutyrate (PBA, 2mM), TUDCA (TUD, 100 µM), and salubrinal (Sal, 30 µM) for 24 hrs before GABA uptake experiment. In A, N=6 batches of cells; in B, N=5-6 batches of new cells. (**C**). Neurons N=5-6 batch of neurons were cotransfected with the wildtype (Wt), the mixture of the wildtype, the GAT-1(A305V) cDNAs (0.25µg:0.25µg) (het). N=5-6 batches of cells. The cells were then treated with vehicle, 4 phenylbutyrate (2mM), TUDCA (100 µM), and Salubrinal (30 µM) for 24 hrs before GABA uptake experiment. (In A to C, ***p < 0.001; ** p < 0.01 vs wt Veh; & p < 0.05, && p < 0.01; &&& vs A305V “Het” Veh. Two-way analysis of variance (ANOVA) and Newman-Keuls test was used to determine significance between different conditions. Values were expressed as mean ± S.E.M).

### Wildtype GAT-1 DNA augmentation rescues SLC6A1 mutations, and combined rescue exceeds either intervention alone

We here first compare the effect on GABA uptake of the mutant GAT-1 in the cells treated with PBA alone or with the wildtype cDNA upregulation at different dosages (Figure 6A-C). We then tested the GABA uptake function in the cells coexpressing the mutant GAT-1(A305V) (AV, 0.25 μg) or GAT-1(S295L) (SL, 0.25 μg) with different cDNA amounts of the wildtype allele (Fig 6A). Augmentation of the wildtype allele cDNA dose-dependently increased the GABA uptake (0.70 for 0.125 μg, 1.1 for 0.25 μg and 1.4 for 0.5 μg for A305V; 0.418 for 0.125 μg, 0.848 for 0.25 μg and 1.11 for 0.5 μg for S295L vs wt=1). We then tested the GABA uptake function in the cells coexpressing the equal amounts of mutant GAT-1 and the wildtype GAT-1 cDNAs and then treated with different concentrations of PBA (0.5 to 4mM) (Fig 6B, C). Cell viability was compromised in the dishes treated with 4mM PBA, which thus was not included. In both A305V (0.65 for veh; 0.81 for PBA 0.5mM; 0.966 for PBA 1mM and 1.18 for PBA 2mM) and S295L (0.49 for veh; 0.65 for PBA 0.5mM; 0.83 for PBA 1mM and 1.04 for PBA 2mM) mutation conditions. PBA (2mM) increased the GABA uptake and reached the maximal GABA uptake functionality. This suggests that the combined approach of PBA plus the wildtype gene augmentation is more favorable than either approach alone. An optimal gene function restoration can be achieved with a lower dose of gene therapy plus PBA treatment.

**Figure 6.**
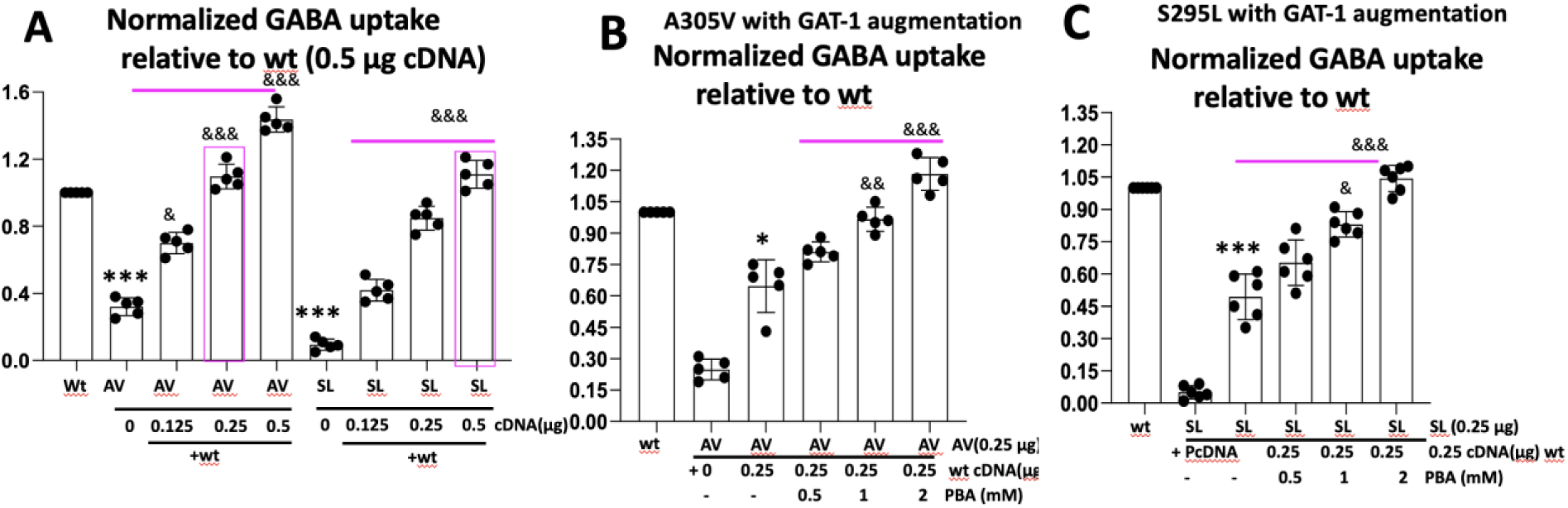
4 phenylbutyrate reduced the dosage of wildtype allele for gene augmentation in mutant GAT-1 transporters in human astrocytes. (**A**) Human astrocytes were transfected with wildtype GAT-1^YFP^ or the mutant GAT-1(A305V)^YFP^ (AV) or GAT-1(S295L)^YFP^ (SL) at the different gene dosage. (**B, C**). Human astrocytes were transfected with wildtype GAT-1^YFP^ or the mutant GAT-1(A305V)^YFP^ (**B**) or GAT-1(S295L)^YFP^ (**C)** with 0.5μg cDNA (0.25 µg of wildtype and 0.25 µg of mutant cDNA) and treated with different amounts of PBA (0, 0.5, 1 and 2mM). In A, B and C, the total cDNA amounts for the wt was 0.5 μg; in the mutant conditions, the total cDNAs were normalized with the vector pcDNA to the total of 0.5 μg. (In A to C, ***p < 0.001; ** p < 0.01 vs wt Veh; & p < 0.05, && p < 0.01; &&& vs AV or SL Veh.

## Discussion

This study identifies SLC6A1 p.Ala305Val as a trafficking-impaired, loss-of-function GAT-1 variant whose dysfunction can be rescued by pharmacologic and genetic interventions acting on distinct but convergent steps in transporter biogenesis. Four findings in this study support this conclusion. First, computational stability predictors uniformly indicate destabilization at Ala305 for both A305V and the residue-matched A305T comparator, with a magnitude consistent with disruption of transmembrane packing. Second, ^3^H-GABA uptake is reduced across HEK293T cells, human iPSC–derived astrocytes, and human iPSC–derived cortical neurons, with A305V exhibiting a GABA deficit like the cells treated with GAT-1 specific pharmacological inhibitors such as Cl-966 and NNC-711. Third, the mutant transporter accumulates in the ER and the total mature transporter pool is reduced, locating the lesion at folding and forward trafficking rather than at substrate binding or surface pharmacology. Fourth, the deficit is reversible: PBA, TUDCA, salubrinal, in addition to our previous studies on BiP and calnexin overexpression, BIX-mediated BiP activation (Deleeuw et al., 2026) and wildtype GAT-1 cDNA augmentation each independently improve function, and the combination of PBA with gene augmentation produces rescue greater than either single intervention in the available experiments.

The mechanistic implication is that SLC6A1–related DEEs, at least for trafficking-impaired missense variants such as A305V and S295L, are not pure dose-haploinsufficiency diseases of the affected gene and not pure pharmacochaperone-responsive misfolding diseases. They are both. Adding more wildtype GAT-1 helps because the absolute pool of transporter is rate-limiting. Adding PBA helps because the fraction of that pool that successfully folds and traffics is rate-limiting. The two strategies therefore act on different segments of the same biogenesis pathway and should be designed to cooperate in a future precision-medicine framework. This framing predicts that pharmacochaperone responsiveness will vary across SLC6A1 variants, with the largest benefit from gene augmentation plus PBA.

The clinical relevance of these findings is direct and the translatability is high. PBA and its commercial form glycerol phenylbutyrate (Ravicti) are already in clinical use for unrelated indications, and an investigator-initiated trial in SLC6A1- and STXBP1-related DEEs is in progress (NCT04937062). The data presented here support continued evaluation of PBA therapeutics in SLC6A1-related disease, and indicate that variant-level functional and trafficking characterization is necessary to guide precision deployment. The data also justifies continued preclinical development of GAT-1 gene-augmentation strategies in parallel with chemical chaperoning, with the expectation that the two modalities will be most effective in combination and can de-risk the potential overdose of gene therapy.

Several limitations of the study should be acknowledged. Although three independent cell systems were interrogated, in vivo confirmation of combined PBA-plus-augmentation rescue in an A305V knock-in mouse model is necessary to establish translational validity. The augmentation experiments use plasmid co-expression as a model of gene therapy which allows us to evaluate the effect of PBA in addition to gene therapy and PBA without the variables and tropism from viral or non-viral delivery in vivo. Finally, the analyses here address two substitutions at a single residue; generalization to the broader SLC6A1 variant landscape will require systematic characterization of pharmacochaperone responsiveness across a representative variant panel, ideally coupled to in-silico stability prediction that has been calibrated against functional readouts.

Despite these limitations, the central conclusion of this study is robust: SLC6A1 A305V causes a well-defined trafficking-linked transporter deficit, that deficit is reversible by interventions acting on protein folding and on transporter gene dose, and the combination provides a rational and translationally accessible framework for precision therapy in SLC6A1-related DEEs. PBA can de-risk gene therapy by maximally leveraging the existing foldable copies of the targeted transporter.

## Conclusions

SLC6A1 p.Ala305Val is a *de novo* missense variant that destabilizes GAT-1, increases endoplasmic reticulum retention, and reduces GABA uptake across heterologous cells and human iPSC–derived astrocytes and cortical neurons. Pharmacologic chaperones (PBA, TUDCA), ER-stress modulators (salubrinal), genetic supplementation of wildtype allele each independently rescue transporter function, and the combination of PBA with gene augmentation produces rescue greater than either alone. These findings define a coherent two-axis therapeutic approach: folding correction plus transporter-dose augmentation—for SLC6A1-related DEEs and motivate variant-level characterization as the next step toward precision medicine.

## Availability of supporting data

Any raw data of functional assay can be made available upon request.

Any clinical information can be made available upon request subject to approval by the appropriate ethical board.

## Authors’ contributions

J.-Q.K. conceived and supervised the study, performed confocal microscopy, analyzed data, and wrote the manuscript. A.J.D performed experiments and wrote the paper. E.G and K.J contributed clinical information. D.S and M.B. performed experiment. J.W. performed AI/structural modeling and stability prediction analyses. All authors reviewed, edited, and approved the final manuscript.

## Acknowledgements

We are grateful to Dr. Wangzhen Shen for her supervision on biochemical assays. The authors thank the family of the affected individual for their participation. We thank the patient communities that have made SLC6A1 research possible. Cell imaging was performed in part through the Vanderbilt University Medical Center Cell Imaging Shared Resource.

